# Micrococcal nuclease sequencing of porcine sperm suggests a nucleosomal involvement on semen quality and early embryo development

**DOI:** 10.1101/2021.03.30.437505

**Authors:** Marta Gòdia, Saher Sue Hammoud, Marina Naval-Sanchez, Inma Ponte, Joan Enric Rodríguez-Gil, Armand Sánchez, Alex Clop

## Abstract

**Background:** The mammalian mature spermatozoon has a unique chromatin structure in which the vast majority of histones are replaced by protamines during spermatogenesis and a small fraction of nucleosomes are retained at specific locations of the genome. The chromatin structure of sperm remains unresolved in most livestock species, including the pig. However, its resolution could provide further light into the identification of the genomic regions related to sperm biology and embryo development and it could also help identifying molecular markers for sperm quality and fertility traits. Here, for the first time in swine, we performed Micrococcal Nuclease coupled with high throughput sequencing on pig sperm and characterized the mono-nucleosomal (MN) and sub-nucleosomal (SN) chromatin fractions.

**Results:** We identified 25,293 and 4,239 peaks in the mono-nucleosomal and sub-nucleosomal fractions, covering 0.3% and 0.02% of the porcine genome, respectively. A cross-species comparison of nucleosome-associated DNAs in sperm revealed positional conservation of the nucleosome retention between human and pig. Gene ontology analysis of the genes mapping nearby the mono-nucleosomal peaks and identification of putative transcription factor binding motifs within the mono-nucleosomal peaks showed enrichment for sperm function and embryo development related processes. We found motif enrichment for the transcription factor Znf263, which in humans was suggested to be a key regulator of the genes with paternal preferential expression during early embryo development. Moreover, we found enriched co-occupancy between the RNAs present in pig sperm and the RNA related to sperm quality, and the mono-nucleosomal peaks. We also found preferential co-location between GWAS hits for semen quality in swine and the mono-nucleosomal sites identified in this study.

**Conclusions:** These results suggest a clear relationship between nucleosome positioning in sperm and sperm and embryo development.

## Background

Current research is starting to evidence that besides the paternal genome, sperm also carries other molecular information to the zygote, including a wide repertoire of RNAs, proteins and epigenetic marks in the form of DNA methylation, retained nucleosomes and histone modifications that can play a role in spermatogenesis, fertility, early embryonic development and even inter- or transgenerational transmission [1–4]. During spermatogenesis, male germ cells undergo a series of profound morphological and functional changes that conclude in mature spermatozoa. In the last stages of spermatogenesis, when the cell is transcriptionally silent, genomic DNA is highly condensed to fit into the sperm head [5] and ensure genomic integrity, fertility and early embryo development [6]. In mammals, the condensation of the spermatozoon chromatin occurs by the progressive replacement of histones by protamines. However, a small fraction of the sperm DNA remains organized in histones as shown (~ 5 to 15%) in humans [4, 7–10], (~ 13.4%) cattle [11] and (~ 1 to 2%) mice [12–14]. These nucleosomes are distributed along the genome following a programmatic pattern with preferential retention at gene promoters and developmental related loci [4, 7, 9, 10]. Nucleosome positioning in the genome and chromatin accessibility are critical in the regulation of gene expression and have been linked to multiple phenotypes [15]. Nucleosomes in sperm may either be leftovers of gene expression during spermatogenesis or also serve as transcriptional instructions upon fertilization for gamete recognition and early embryo development [16]. Thus, this information can be of great value to shed light into the catalog of genomic regions and genes related to sperm biology and early embryo development and thus help identifying elusive molecular markers for such traits.

In livestock, the architecture of the sperm chromatin at the genomic level has only been investigated in cattle [11]. In pigs (*Sus scrofa*), we and others have studied the sperm RNA [17–19] and protein populations [20] as well as the DNA methylation patterns [21] but the sperm’s chromatin structure remains unexplored.

In this study, we profiled for the first time in pigs, the nucleosome location in the sperm’s chromatin of sperm. Pig spermatozoa were digested with micrococcal nuclease (MNase) and the resulting nucleosome-associated DNAs were subjected to high throughput sequencing. We characterized the mono-nucleosomal (MN) and sub-nucleosomal (SN) chromatin fractions and assessed their correlation with sperm RNA levels and their co-location with genomic regions associated to semen traits.

## Methods

### Sample collection

Ejaculates from two healthy Pietrain boars, replicate A (16 months of age) and replicate B (9 months old), from two artificial insemination studs were obtained by specialists during their routine sample collection using the gloved hand method [22] and were immediately diluted (1:2) in commercial extender. Both ejaculates showed high semen quality. Semen samples were purified and processed as in [23], and stored at −80°C until further use.

### Micrococcal nuclease digestion, library construction and sequencing

Chromatin digestion was performed as previously described [4] with minor modifications. 40 million spermatozoa cells were incubated with 5 U of MNase (Sigma-Aldrich) for 7 min at 37°C. The resulting digested DNA was evaluated in a 1.5% agarose 1 x TAE gel electrophoresis. The nuclease digested DNA (which included both the MN and SN fractions) was directly used for library prep after purification with the Agencourt AMPure XP beads (Beckman Coulter). The MN and SN fractions were separated after sequencing and read mapping using bioinformatics tools as described below. Purified DNA was subjected to quality control including quantification with the Qubit™ DNA HS Assay kit (Invitrogen) and Nanodrop (Thermo Scientific Fisher) as well as size and concentration assessment with a 2100 Bioanalyzer and the Agilent DNA 1000 Kit (Agilent Technologies). We also extracted gDNA from the two sperm samples as described by Hammoud and colleagues [4]. The two gDNAs were pooled to be used as input. Pooled gDNA were sheared to obtain 100 bp long fragments using a Covaris S2 instrument (Covaris Inc), according to the manufacturer’s instructions. Sequencing libraries of the two MNase treated samples and the input gDNAs were prepared with the PrepX™ DNA Library Kit (Takara). The libraries were used as a template to generate 50 bp paired end (PE) reads in an Illumina HiSeq2500 system.

### MNase-seq data preprocessing and quality evaluation

Raw sequencing data was filtered to remove low quality bases, adaptor sequences and short reads (< 25 bp length) with Trimmomatic v.0.36 [24]. Trimmed reads were aligned to the porcine genome (Sscrofa11.1) with Bowtie2 v.2.4.1 [25] fitting default parameters except “--very sensitive”. Pearson correlation of the read coverages for genomic regions along the genome was calculated using the multiBamSummary file based on the two MNase samples with the tool deepTools v.3.3.2 [26] with options: “plotCorrelation --removeOutliers –skipZeros”. Since the correlation between the samples was very high, the reads from the two replicates (MN_1: replicate A; MN_2: replicate B) were merged and processed as a pool with the aim to increase sequencing depth and peak detection power and accuracy. Then, the MN and SN fractions were bioinformatically separated based on the genomic distance between the two paired reads. To extract the reads from the SN fraction, with a length slightly below 100 bp (see Figure 1a), we used the parameter “--maxins 110”, which indicates that the mapped paired-end reads should not exceed 110 bp. To obtain the reads from the MN fraction, with a length around 150 bp, we used the parameters “--minins 111” and “--maxins 1000”. Subsequently, duplicate reads of these fractions and the input sample were removed with Picard-Tools MarkDuplicates v.1.56 (http://picard.sourceforge.net). Finally, for both fractions, peak calling was performed with MACS2 v.2.1.0 [27] with “-q 0.05 -B -g hs” and compared against the input sample. Visualization of the genomic heatmaps from the MNase-seq signals using transcription start sites (TSS) as reference points was carried with the deepTools [26] computeMatrix tool with parameters: “reference-point -b 500 -a 500 –skipZeros” and plotHeatmap tool using “--refPointLabel ‘TSS’“.

**Figure 1.**
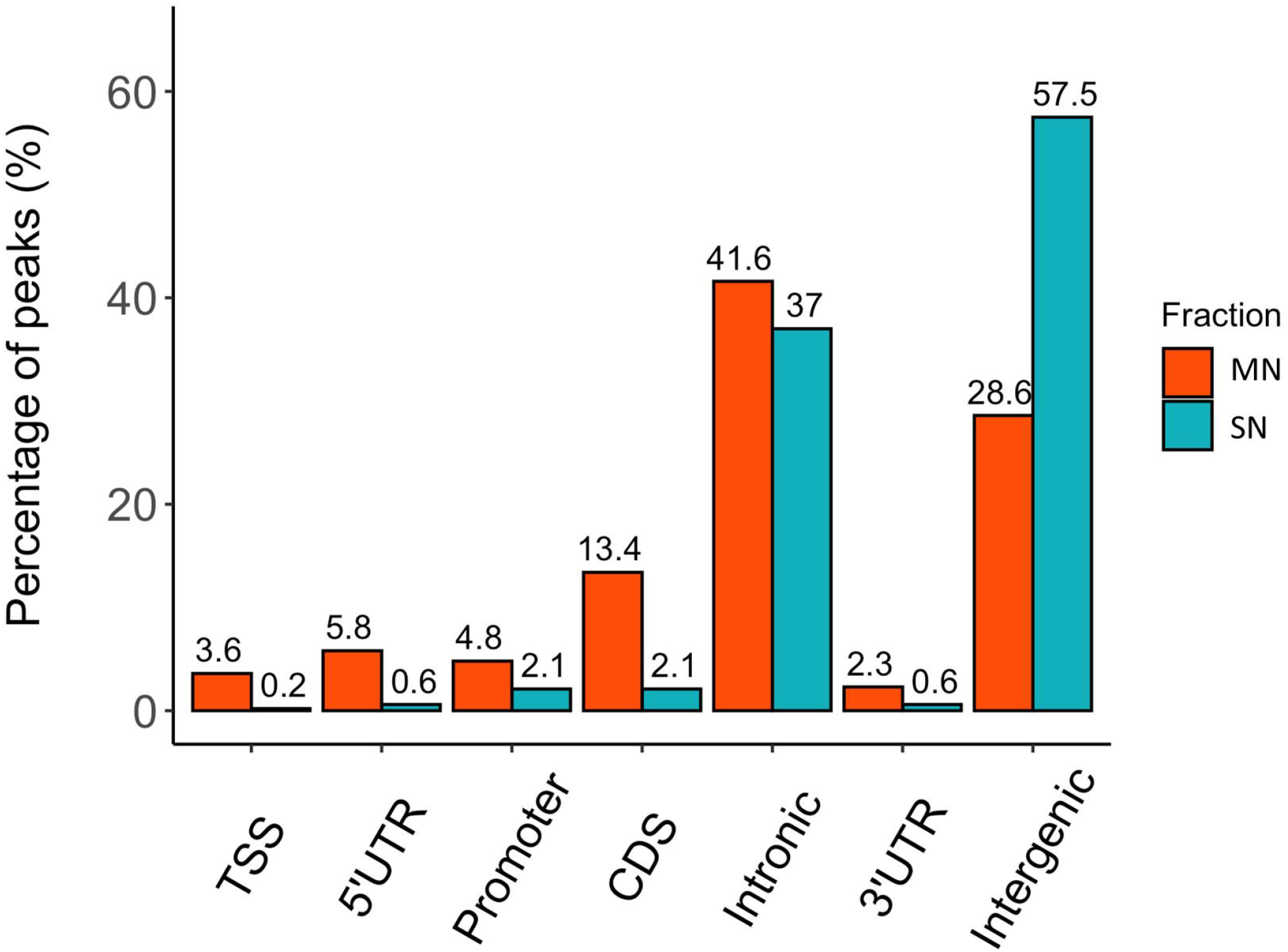
Distribution of the MN and SN peaks relative to gene features in the pig genome. Peaks were classified according to their co-location with gene features as, Transcription Start Site (TSS), 5’ untranslated region (5’UTR), 3’UTR, promoter, coding sequence (CDS), intronic and intergenic.

### Peak location relative to gene annotation

Peaks were categorized by their position in relation to gene features as annotated in Ensembl (v96) using BEDtools v.2.29.2 [28] intersect. The peaks were classified as mapping to TSSs, promoters, overlapping 5’ untranslated region (UTR), 3’ UTR, exonic, intronic and intergenic as carried in [29].

In order to determine any potential positional preference of the peaks within the gene features, 100 iterative permutations of the peak genomic locations with the same number of peaks following the same peak width distribution but with randomized genomic locations were carried using BEDtools [28] shuffle. Subsequently, the randomized peaks were assigned to genomic features according to the genomic position within these features. These results were then contrasted against our results on real data with a one-sample T-test (one-tailed).

### Gene Ontology analysis of the genes near MNase peaks

The genes mapping within or less than 5 kbp apart from the identified peaks were extracted from Ensembl v96 with BEDtools [28] closestBed and were used for Gene Ontology (GO) enrichment analysis. GO was carried out with Cytoscape v.3.8.2 plugin ClueGO v.2.5.7 [30] using the Cytoscape’s porcine dataset and default settings. Only the significant Bonferroni corrected *p*-values were considered.

### Motif enrichment analysis at the MN and SN underlying sequences

Motif enrichment analysis was carried out with the software HOMER v.4.10.0 [31] findMotifsGenome.pl function with default parameters.

### Positional conservation of chromatin-associated DNA with human sperm

A comparable human sperm MNase-seq dataset (GSE15701) from Hammoud et al. [4], obtained from a pool of four donors was downloaded from the Gene Expression Omnibus database. The genomic coordinates from the human sperm MN peaks were liftover to Sscrofa11.1 using the UCSC liftover tool [32] with default parameters except for “-minMatch=0.1”. For the enrichment evaluation, the genomic location of liftover peaks were randomized 100 times using BEDtools [28] shuffle and overlapped to the porcine MN peaks identified in this study using BEDtools [28] intersect. Thereafter, the results were contrasted to the real liftover genomic data with a one-sample T-test (one-tailed).

### Integration with other -omics data

Although mature sperm is transcriptionally inactive, RNAs in sperm may both reflect preceding events in transcriptionally active spermatogenic precursor cells during male germ cell development and have a role in early development after fertilization [2, 33]. Thus, we hypothesized that a relationship between the location of retained nucleosomes in sperm and the sperm transcriptome profile may exist. To test this hypothesis, sperm transcriptome data from 40 Pietrain boars, including 40 total and 35 small RNA-seq datasets previously generated by our group (NCBI’s BioProject PRJNA520978) was used. These datasets provided a list of 4,120 protein coding RNAs [34], 1,574 circular RNAs (circRNAs) mapping in pig chromosomes [18] and 6,729 PIWI-interacting RNAs (piRNAs) [19]. The averaged RNA abundances from all the samples were used for further analysis. Protein coding genes were classified according to their RNA abundances measured in Fragments per Kilobase per Million mapped reads (FPKM) as (i) absent (< 1 FPKM); (ii) low abundance (≥1 to <10 FPKM); (ii) intermediate abundance (≥ 10 to < 100 FPKM) and (iii) high abundance (≥ 100 FPKM). Then, the existence of positional co-location enrichment between each of these gene fractions and the MNase peaks was assessed using the Fisher Exact Test (two-tailed). We considered as co-location any distance below 5 kb between a RNA and a MN or SN peak. Moreover, we compared the RNA abundance of the genes that co-located with MN and SN peaks with the RNA levels of the whole set of genes annotated in the pig genome employing the Kruskal-Wallis test using RNA abundances stabilized with the log2 transformation.

We also studied the positional enrichment of circRNAs or piRNAs at MNase peaks. First, the genomic regions of the circRNAs or piRNAs were iteratively randomized 100 times using BEDtools [28] shuffle. Then, the enrichment of the real piRNA and circRNA genomic coordinates at MNase peaks was assessed by comparison with the overlap of the randomized locations with the MNase peaks using a one-sample T-test (one-tailed).

MNase profiles were also evaluated against the genomic regions showing genetic association with sperm quality traits in Pietrain in a Genome-Wide Association Study (GWAS) carried by our group and which included the two samples used in this study [34]. This GWAS provided 18 GWAS regions associated to the percentage of sperm cells with head abnormalities (7 hits), neck abnormalities (4), percentage of abnormal acrosomes after 5 minutes incubation at 37°C (2), the ratio of abnormal acrosomes after 5 minutes and 90 minutes of incubation (1), percentage of motile spermatozoa after 5 minutes incubation (2) and after 90 minutes incubation (2, one hit shared with motility after 5 minutes incubation) and the percentage of sperm cells with proximal droplets (1). These regions spanned in total, 16.5 Mbp (0.7%) of the porcine autosomal genome [34]. Again, to evaluate whether MNase peaks were enriched at GWAS hits, the genomic regions of the GWAS hits were iteratively randomized 100 times using BEDtools [28] shuffle. This randomized overlap was compared to the overlap of the real data using the one-sample T-test (one-tailed).

## Results

### MNase digestion, library preparation and data preprocessing

Nucleosome-associated DNAs were released from boar sperm using an optimized MNase digestion protocol. Echoing the results observed in human [4, 10, 11] and cattle [11] spermatozoa, the MNase treatment generated ~147 bp fragments corresponding to the MN fraction and an additional ~ 100 bp fragment corresponding to the SN fraction (Additional file 1).

The sequencing of the libraries yielded more than 43 million PE reads for each MNase biological replicate and 41.8 million PE reads for the input sample (Table 1). In average, 90.2% of the MNase reads and 92.7% for the input reads mapped to the porcine genome (Sscrofa11.1). As the genome-wide profiles of MNase sensitivity of the two samples showed high correlation (Pearson R = 0.87) (Additional file 2), the mapped reads from the two samples were merged and further analyzed as a pool with the aim to increase read depth and peak call sensitivity and accuracy. The final number of PE reads in each, the MN and SN fractions, was 62.1 million and 7.1 million, respectively (Table 1).

**Table 1.**
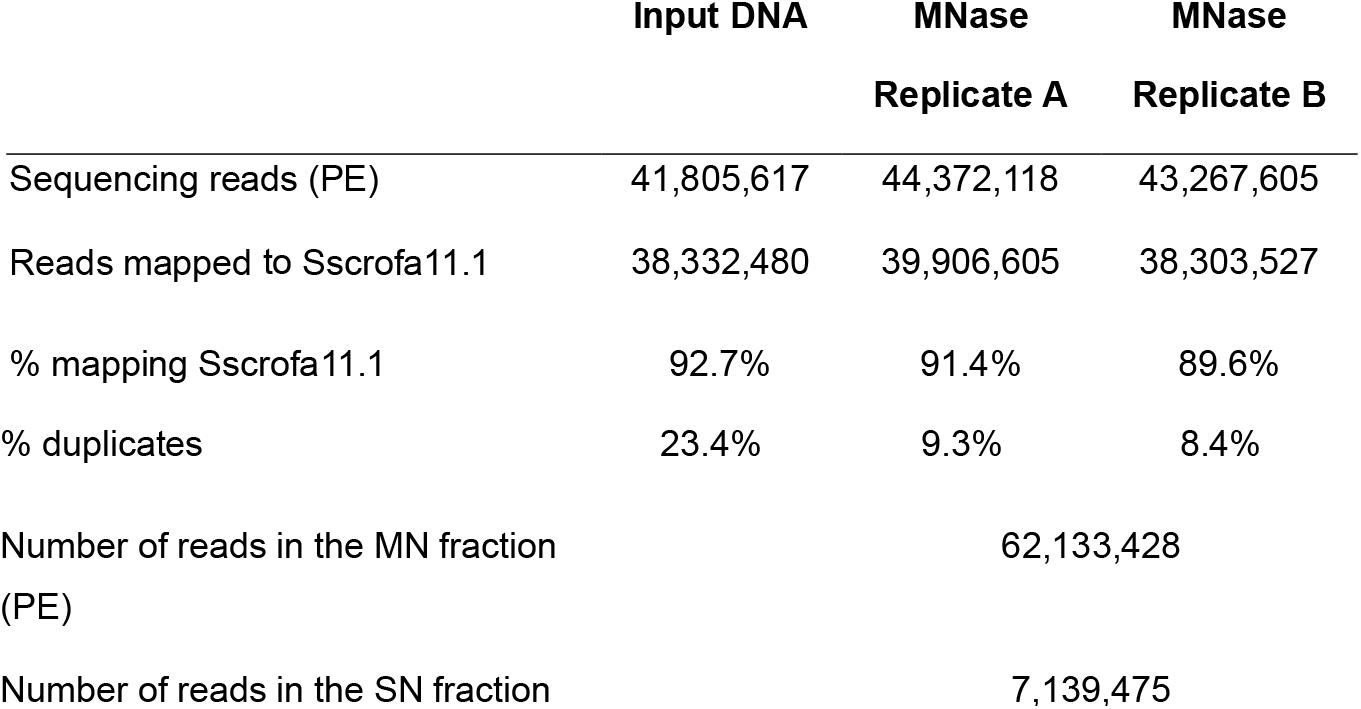
MNase-seq sequencing and pre-processing metrics for the input and the two MNase biological replicates.

|  | Input DNA | MNase<br>Replicate A | MNase<br>Replicate B |
| --- | --- | --- | --- |
| Sequencing reads (PE) | 41,805,617 | 44,372,118 | 43,267,605 |
| Reads mapped to Sscrofa11.1 | 38,332,480 | 39,906,605 | 38,303,527 |
| % mapping Sscrofa11.1 | 92.7% | 91.4% | 89.6% |
| % duplicates | 23.4% | 9.3% | 8.4% |

|  |  |
| --- | --- |
| Number of reads in the MN fraction<br>(PE) | 62,133,428 |
| Number of reads in the SN fraction<br>(PE) | 7,139,475 |

#### Global characterization of the MN and SN peaks

We identified 25,293 MN and 4,239 SN peaks (Additional file 3 and 4). The MN peaks averaged 270 bp in width and covered 0.3% of the porcine genome. The SN peaks were 141 bp wide and covered 0.02% of the genome. Most MN (70.2%) and SN (94.5%) peaks were annotated in intronic and intergenic regions (Figure 1). MN peaks were significantly enriched near TSS (*p*-value = 6.4e-192) (Figure 2). This result was not replicated in the SN peaks (*p*-value = 0.20). A total of 9,128 and 1,688 genes overlapped or localized less than 5 kbp apart from a MN and a SN peak, respectively (Additional file 5). The two MN and SN fractions co-existed in 1,145 genes (Additional file 5), which corresponds to 75% of the SN-associated genes.

**Figure 2.**
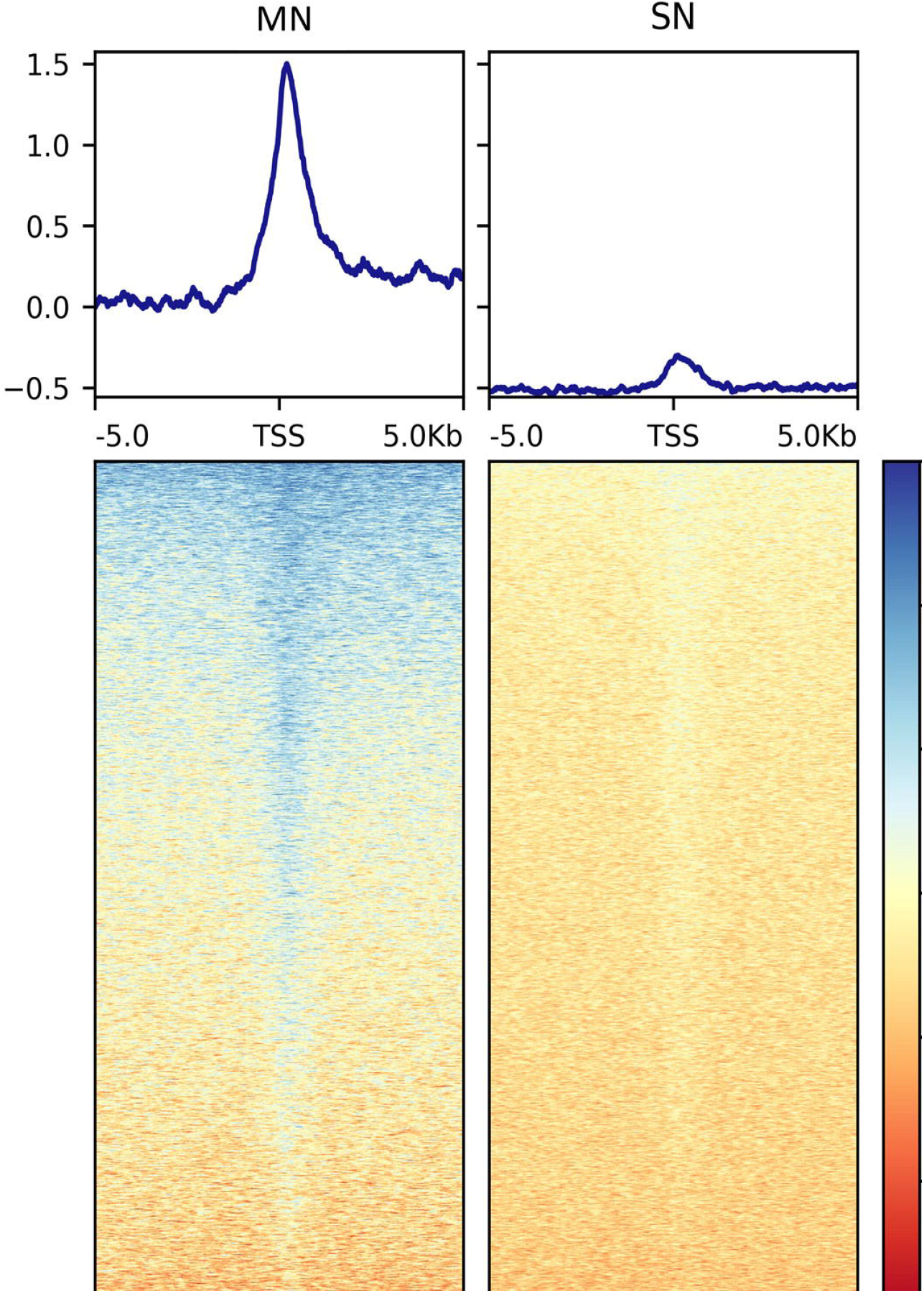
Genomic heatmaps depicting the normalized MNase-seq signal centered at TSS for the MN and the SN peaks. The x axis shows the genomic location relative to the TSS. The y axis indicates the MNase-seq signal intensity.

To interrogate the potential function of the nucleosome associated DNA, we carried out GO analysis of the genes overlapping or mapping less than 5 kb away from these peaks. For the MN peaks, the most significantly enriched terms were related to chemical perception, development (including the nervous system, multicellular organism, tissue, embryo, etc) and cell and organ morphogenesis (Additional file 6). The genes located in the SN fractions were enriched for less GO terms and showed weaker significances and included nervous system process, cell projection, regulation of GTPAse activity, organelle organization and Golgi vesicle transport (Additional file 7). To delve further into the potential functions of the nucleosome retention in sperm, the genomic sequences underlying the peak regions were subjected to sequence motif enrichment analysis. In MN peaks, we identified enrichment for motifs from 55 plant and 57 animal (mostly mammalian: mouse and human) transcription factors (TFs) (Additional file 8). The mammalian included 15 TFs from the bHLH class (MyoD, MyoG, Myf5, NeuroG2, Olig2, Tcf21, Twist2, etc) and other TFs related to embryo development and implantation (Atf4, CHOP, Erra, EBF1, E2F2, Foxh1, HOXA1, HOXA2, MAFK, Pax8, IRF1, IRF3, PR, Rfx1, RORg, RUNX2, Zic, Zic3). They also included TFs related to spermatogenesis or sperm function which have also been related to embryo development such as CUX1, CTCFL, PAX5 and Smad2. Finally, this list also contained Znf263 and FOXA1, two TFs that provide a direct link between the paternal gamete and the embryo (Additional file 8). One study identified 514 genes with paternal preferential expression during human early embryo development and proposed *ZNF263* as the strongest candidate in regulating the expression of these genes [35]. We identified the pig genes that map less than 500 bp apart from a MN peak harboring a Znf263 motif and searched the overlap with the 514 human genes showing paternal preferential expression in the early embryo. 3,264 genes mapped less than 500 bp away from a MN peak with Znf263 binding motif in pig sperm. These corresponded to 3,288 human orthologs, 92 of which (18% of 514) also showed preferential paternal expression during human early embryo development [35] (Additional file 9). This concordance nearly reached statistical significance with the hypergeometric test (*p*-value = 0.052). The SN peaks presented enriched motifs for a similar catalogue of TFs identified in the MN fraction (Additional file 10).

A comparable human sperm MNase-seq dataset, which included pool data from four donors [4] was employed to assess the inter-species conservation of the genomic location of the MNase peaks. Results in human were similar to these detected in our analysis in pig. 25,122 peaks were annotated in the human MN fraction, 20,641 of which were successfully converted to the pig genome coordinates. 27% of the 5,540 human nucleosome associated sites overlapped with our porcine MN peaks, thus showing that the nucleosomes in human and pig sperm tend to co-locate (*p*-value = 4.8e-258).

#### Functional characterization of the nucleosome-associated DNA

To identify positional relationships between the nucleosome-associated regions and transcriptional activity, the locations of the MN and SN peaks were compared with the repertoire of RNAs that our group identified in porcine sperm [34]. 12,125 protein coding genes were absent in sperm. On the contrary, 5,814, 3,521 and 598 genes were classified as being at low, moderate and high abundance in sperm, respectively. The location of the genes that are present in spermatozoa was, regardless of their abundance, highly enriched (low abundance: *p*-value = 1.8e-72, moderate abundance: *p*-value = 1.5e-91, high abundance: *p*-value = 1.1e-11) at the MN peaks when compared to the catalog of absent genes (Table 2). Similarly, the SN peaks were also enriched for the low (*p*-value = 5.0e-17), moderate (*p*-value = 3.6e-47) and highly abundant (*p*-value = 3.0e-5) genes, when compared to the set of absent genes (Table 2). In line with these results, the average RNA abundance of the genes mapping to both MN (*p*-value = 4.9e-19) and SN (*p*-value = 4.3e-22) peaks was significantly higher than the average abundance of all the genes annotated in the pig genome (Figure 3).

**Table 2.** Distribution of the protein coding genes within the MN and SN fractions, according to their RNA abundance in sperm.

|  |  | Genes within the MN peaks |  |  | Genes within the SN peaks |  |  |
| --- | --- | --- | --- | --- | --- | --- | --- |
|  |  | Within MN peaks | Not present | *p*-value | Within SN peaks | Not present | *p*-value |
| Gene' s RNA levels in sperm | Not present | 4083 | 8042 |  | 644 | 11481 |  |
|  | Low abundance | 2774 | 3040 | 1.80E-72 | 504 | 5310 | 5.00E-17 |
|  | Intermediate abundance | 1857 | 1664 | 1.50E-91 | 455 | 3066 | 3.60E-47 |
|  | High abundance | 296 | 302 | 1.10E-11 | 60 | 538 | 3.00E-05 |

**Figure 3.**
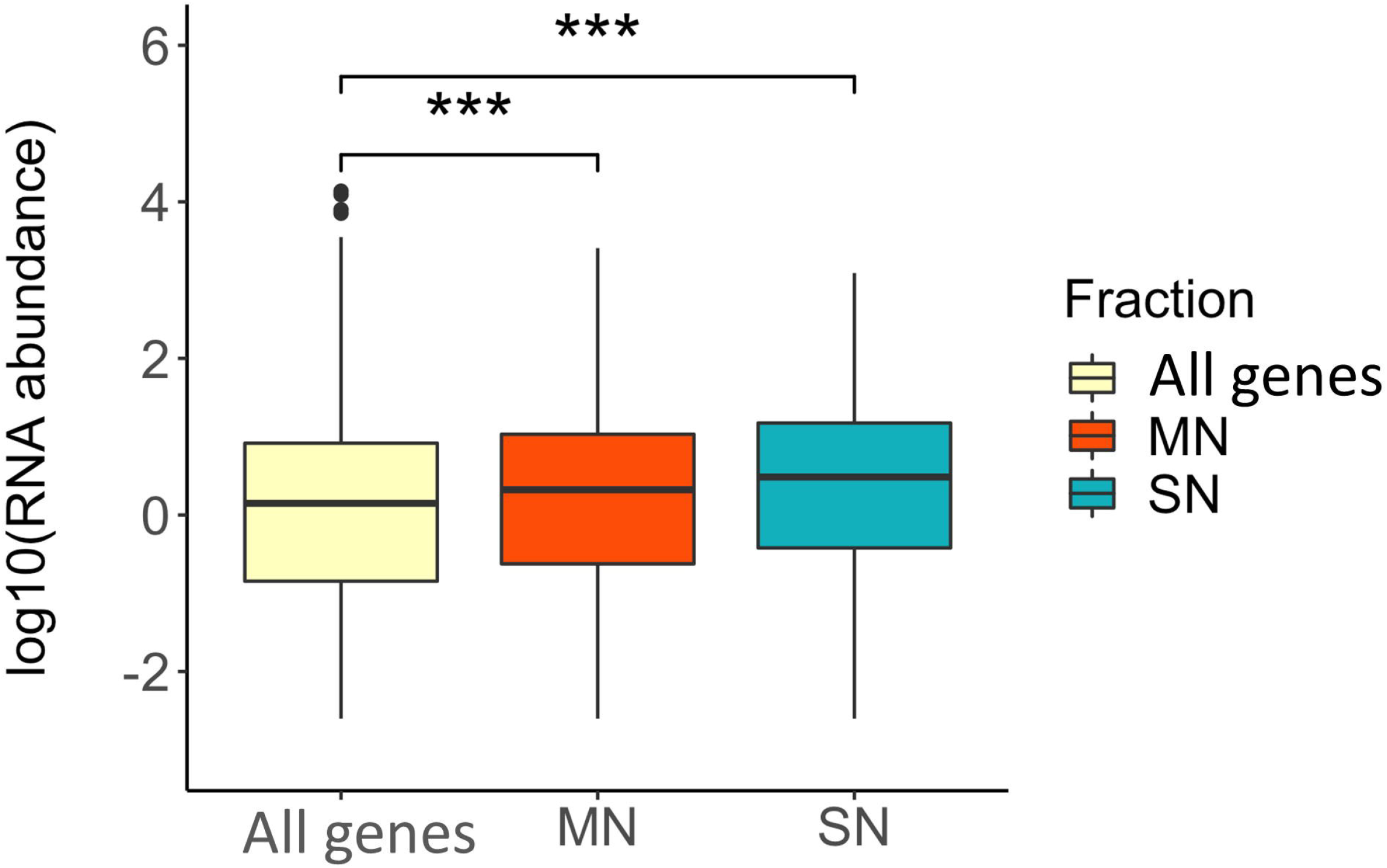
Box plots showing the average RNA abundance of (i) all genes present in the genome (yellow), (ii) the genes present in the MN fraction (red) and (iii) the genes present in the SN fraction (blue). ***: *p*-value ≤ 0.001.

A similar enrichment at MN peaks was also observed for the catalog of 1,574 sperm circRNAs [18]. They overlapped to 3,198 MN peaks and showed an enrichment when compared with the results of the randomized analysis (*p*-value = 2.0e-172). Likewise, 148 sperm circRNAs associated to sperm motility in swine overlapped to 14 MN peaks and showed preferential location within nucleosomal sites (*p*-value = 7.1e-64, overlapping to 14 MN peaks). Recently, a population of 6,729 sperm piRNAs have been annotated in pig [19]. The piRNA regions were significantly mapping more frequently within MN peaks (*p*-value = 4.8e-53, overlapping to 65 MN peaks). This enriched co-location was also observed (*p*-value = 5.6e-32, overlapping to 18 peaks) for the set of piRNA genomic regions (involving 1,355 piRNAs) which abundance in sperm correlated with different sperm phenotypes including the percentage of motile sperm, the percentage of morphological abnormalities or the percentage of viable cells [19].

Our group recently published a GWAS for sperm quality traits in Pietrain boars [34] which we used to evaluate the co-location of MNase sites with regions associated to these traits and establish potential links between nucleosome retention and sperm quality. We found a significant co-location (*p*-value = 9.3e-108) between both features, with 189 MN peaks overlapping to a GWAS hit. This trend was also observed between GWAS regions and SN peaks (*p*-value: 5.3e-49) with 24 SN peaks overlapping a GWAS hit. These co-locations involved mostly regions associated to the percentage of head and neck abnormalities and, to a lesser extent, to the percentage of abnormal acrosomes and sperm motility.

### Discussion

In this study we describe, for the first time, the genome-wide characterization of MNase digested chromatin - indicating the genomic location of retained histones - in porcine spermatozoa. Echoing the results observed in humans [4, 10, 11], the digestion resulted in two DNA bands (Additional file 1). One band corresponded to the MN fraction including segments of the genome harboring one nucleosome, which typically consists of ~ 147 bp of DNA wrapped around the histone octamer comprising two copies of the core histones H2A, H2B, H3, and H4 [36]. A smaller SN band of ~ 100 bp was also identified. A SN band has been also detected in yeast [37], *Xenopus laevis* sperm [38] and human sperm [10] and may correspond to partially unwrapped nucleosomes that have lost one of the H2A and H2B dimers. Proteomic analysis of SN bands in *Xenopus* revealed association with chromatin regulatory proteins (H3, H4, H1FX, CBX3, WDR5) [38]. Other articles describe that these remodeling complexes can facilitate TF binding to SN when this includes their binding sequence. Then, the presence of SN constitutes a novel physical signature of active chromatin [39].

The extraction of nucleosomal DNA was less efficient than in experiments carried in the sperm of other species [4, 9–11, 13], and it did not yield sufficient amount of DNA from the electrophoretic bands for high throughput sequencing. Nevertheless, the total amount of digested DNA was enough for sequencing and fragment size separation after read mapping using bioinformatics tools. To directly sequence the MN and SN fractions, future experiments will require processing a larger number of sperm cells to isolate sufficient DNA from each MNase electrophoretic band.

The nucleosome-associated DNAs covered ~ 0.3% of the porcine sperm, which is nearly ten-fold and 75% more nucleosome retention than what has been described in human [4, 7–10] and mouse [12–14], respectively. Although this difference could in part have technical causes, we cannot rule out interspecies variability, which could be partially driven by differences in the protamine amino acid content resulting in changes on the extent or arrangement of the protamine disulfide bonding [40].

An increasing body of evidence in other animal species points towards a programmatic nucleosome retention in the sperm genome [4, 10]. Most MN (70.2%) and SN (94.5%) peaks were annotated in intronic and intergenic regions (Figure 1), which is not unexpected since these gene features cover the vast majority of the genomic space. As a matter of fact, MN locations were enriched near TSS sites (Figure 2), providing the first indication that nucleosome retention in sperm is not stochastic and may relate to genomic functional activity. This hypothesis is further supported by the results we obtained on the comparison of the genomic location of the MN and SN peaks in porcine sperm and in other species; the gene annotation of the pig genome; the prediction of TF binding motifs at MN sites and our own data on the pig sperm transcriptome and GWAS for pig semen traits.

In concordance with previous findings in other species, our results in swine showed that nucleosome peaks were predominantly retained in gene promoters and significantly overlapped with their human synthenic regions [4], thereby indicating a degree of inter-species conservation in gene regulation, normally associated to the maintenance of important functions. Nonetheless, no enriched overlap was observed when comparing our results in pigs with the MNase data obtained in cattle sperm [11] with only 9 and 19 overlapping peaks with the two bovine samples (*p-*value > 0.05). This discrepancy might be attributed to either technical differences in the protocol or to a suboptimal annotation of the cattle genome when compared to human as indicated by the limited success in the conversion of genomic coordinates from the cattle to the pig genomes (only 45% and 49% of the 2,256 and 8,446 peaks called in the two cattle samples were successfully converted to pig genomic coordinates).

We observed a larger number of strongly enriched GO terms for the catalog of genes mapping nearby the MN when compared to the SN fractions (Additional file 6 and 7). The most enriched terms for the MN fraction included sensory perception, which is important for sperm homeostasis (e.g., *SOD2;* [41]), capacitation (e.g., *TRPV1;* [42]) and chemotaxis guidance to the egg for fertilization (e.g., *CA6;* [43]) and cell differentiation, development and morphogenesis, all with obvious links to embryogenesis (Additional file 6).

The search for TF motifs at MN peaks also indicated that MN peaks may carry instructions for embryo development. First, we identified enriched motifs for several TFs related to embryo development (Additional file 8). Some of these genes clearly point toward a role of the paternal gamete in regulating embryo development. This is the case for *ATF4* [44] and *CHOP* [45], which are relevant in the stress response of the mouse embryo at key embryogenesis steps such as zygotic genome activation. *FOXA1* is bound, in the chromatin of human sperm, to genomic regions corresponding to enhancer elements in embryonic stem cells and in several cell types of the embryo, thereby suggesting that the position of this TF in sperm guides the location of relevant enhancers for embryo development [16]. The identification of Znf263 motifs is another relevant finding. *ZNF263* has been proposed as a master regulator of the genes showing paternal preferential expression in the early developing human embryo [35]. Interestingly, we found a nearly significant concordance between the genes with paternal preferential expression in the human embryo and the pig orthologs mapping near a MN peak with predicted ZNF263 binding motif in sperm (Additional file 9). This data suggests that nucleosome retention in sperm has guiding relevance in the early embryo development.

Although mature sperm is transcriptionally silent, it carries a wide repertoire of RNAs that are implicated in spermatogenesis, fertilization, embryo development and offspring phenotype [reviewed in: 2], a large proportion of which were transcribed in transcriptionally active spermatogenic cells during previous stages of spermatogenesis. Our group has generated sperm RNA-seq [18, 19, 34] and GWAS data for semen quality traits [34] on a set of Pietrain boars. The 2 boars used for the MNase study, also Pietrain, were included in the GWAS. The boars used for RNA-seq, also used in the GWAS, did not include the 2 MNase samples. The regulation of transcription is modulated by nucleosome occupancy and the accessibility of the genome to the transcriptional machinery [46]. Our results show that the genes mapping within or nearby retained nucleosomes tend to display larger RNA abundance than the full set of genes annotated in the porcine genome (Figure 3). These results also indicate that the genes present in sperm tend to map to nucleosome-retained loci in both the MN and SN fractions. Not only protein coding but also regulatory RNAs (circRNAs and piRNAs), were also significantly enriched at MN peaks. Again, this data supports the notion that nucleosome occupancy is key in modulating gene expression. Moreover, the enriched co-location between the MN and SN sites and the catalog of circRNAs [18] and piRNA regions [19] which abundance correlated with sperm quality phenotypes further suggests that this regulation during spermatogenesis may have phenotypic consequences on semen quality and reproduction. Thus, the nucleosomes and sub-nucleosomes associated to these RNAs may be leftovers from spermatogenesis and might provide useful information for the identification of markers of abnormal spermatogenesis and sperm quality.

In light of these results, and since nucleosome positioning can modulate gene expression [47–49], we hypothesized that sperm-retained nucleosomes could encompass DNA variants altering elements such as promoters or enhancers regulating the expression of genes that played a role during of spermatogenesis. In such case, we should observe some degree of co-location between MNase peaks and GWAS hits. Indeed, we observed this trend between the MN and SN peaks with the GWAS hits for semen quality [34]. This result further supports the hypothesis that nucleosome positioning in sperm is not stochastic and may be related to the expression of key genes during previous stages of spermatogenesis that are important for semen biology. The preference for GWAS hits for the percentage of head and neck abnormalities is most probably due to the fact that these were the traits that showed more hits (11 of 18) in our GWAS [34] and does not seem to indicate a real preferential relationship between MN positioning and sperm morphology.

### Conclusion

In conclusion, we can speculate that nucleosome retention in mature spermatozoa is related to transcriptional regulation during spermatogenesis and that it is also an instructional contributor to early embryo development. Hence, interrogating the nucleosome occupancy in the sperm chromatin could help elucidating the biological basis of sperm quality and early embryo development and could assist in the search for biomarkers for these sets of traits.

## List of abbreviations

cirRNA: circular RNA
FPKM: Fragment per Kilobase per Million mapped reads
GWAS: Genome-Wide Association Study
MNase: Micrococcal nuclease
MN: Mono-Nucleosomal
piRNAs: Piwi-interacting RNAs
SN: Sub-Nucleosomal
TSS: Transcription Start Site
TF: Transcription Factor

## Declarations

### Ethics approval and consent to participate

The ejaculates obtained from pigs were privately owned for non-research purposes. The owners provided consent for the use of these samples for research. Specialized professionals at the farm obtained all the ejaculates following standard routine monitoring procedures and relevant guidelines. No animal experiment has been performed in the scope of this research.

### Consent for publication

Not applicable.

### Availability of data and material

We are now in the process of submitting the fastq files relative to replicate A, replicate B and the input pool to NCBI’s SRA.

### Authors’ contributions

MG, AS and ACl conceived and designed the experiment; JR-G carried the phenotypic analysis; MG carried the laboratory work with support from IP and SSH; MG made the bioinformatics and statistical analysis; MN provided bioinformatics support. MG analyzed the data, with special input from SSH and ACl. MG and ACl wrote the manuscript; all authors discussed the data and read and approved the contents of the manuscript.

## Acknowledgements

We would like to thank all the members of Hammoud lab for their support.

## Additional files

### Additional file 1 (PDF)

DNA binding pattern of Pig spermatozoa chromatin digested with Micrococcal nuclease.

**a.** Agarose gel electrophoresis of the sperm chromatin after Micrococcal nuclease digestion resulted in a mono-nucleosomal (MN) (~147 bp) and a sub-nucleosomal (SN) (<100 bp) bands. Left: 100 bp DNA ladder; Right: ~ 300 ng of MNase digested sperm chromatin. **b.** Bioanalyzer electropherogram profile of a MNase digested sample showing the SN and MN DNA fractions. *x*-axis: bp fragment length; *y*-axis: FU (signal intensity) of the DNA fragments.

### Additional file 2 (PDF)

Correlation between MNase-seq signals between the two biological replicates.

Scatterplots showing the Pearson’s correlation of the normalized MNase-seq signals between the two biological replicates.

### Additional file 3 (XLSX)

List of the mono-nucleosomal peaks in pig sperm.

List of the MN peaks detected in porcine sperm. For each peak, the list shows, chromosome, start position, end position, width, -log10(*p*-value) and fold enrichment.

### Additional file 4 (XLSX)

List of sub-nucleosomal peaks in pig sperm.

List of the SN peaks detected in porcine sperm. For each peak, data includes chromosome, start position, end position, width, -log10(*p*-value) and fold enrichment.

### Additional file 5 (XLSX)

List of Ensembl genes annotated in the pig genome mapping less than 5 kb away from a MNase peak. MN: gene mapping less than 5 kb from an MN peak. SN: gene mapping less than 5 kb away from a SN peak. MN+SN: gene mapping less than 5 kb from a MN and a SN peak.

### Additional file 6 (XLSX)

List of biological processes from the Gene Ontology analysis of the genes mapping less than 5kb away from a MN peak.

GO biological process terms with significant Bonferroni corrected *p*-values (*p*-val ≤ 0.05) and their associated genes.

### Additional file 7 (XLSX)

List of biological processes from the Gene Ontology analysis of the genes mapping less than 5kb away from an SN peak.

GO biological process terms with significant Bonferroni corrected *p*-values (*p*-val ≤ 0.05) and their associated genes.

### Additional file 8 (XLSX)

List of sequence motifs enriched within the mono-nucleosomal peaks.

Motif name, *p*-value, percentage of targets sequences with motif, comments on the cognate transcription factor. Comments on the cognate transcription factor contains comments for these transcription factors with direct relevant functions in spermatogenesis or embryo development mostly obtained from searches in the GeneCards^®^ database (https://www.genecards.org) and PubMed. Some of these binding motifs are from studies in Arabidopsis, yeast, Drosophila or *C. elegans* and were not included in the searches.

### Additional file 9 (XLSX)

List of the genes mapping less than 500 bp from a MN peak harbouring a predicted Znf263 binding motif for which the human ortholog have paternal preferential expression in early embryo.

The list of human orthologous genes showing paternal preferential expression in human early embryos was obtained from the article “Single-Cell Transcriptome Analysis of Uniparental Embryos Reveals Parent-of-Origin Effects on Human Preimplantation Development” (Cell Stem Cell 2019, 25:697–712).

Cells in clear green correspond to data from the parent-of-origin RNA expression in human embryo (genes with paternal preferential expression at the 8-cell and morula states embryo). Cells in clear grey indicate data from our study on genes mapping less than 500 bp away from MN peaks with a predicted Znf263 binding motif.

### Additional file 10 (XLSX)

List of sequence motifs enriched within the sub-nucleosomal peaks.

Motif name, *p*-value, percentage of targets sequences with motif, comments on the cognate transcription factor. Comments on the cognate transcription factor contains comments for these transcription factors with direct relevant functions in spermatogenesis or embryo development mostly obtained from searches in the GeneCards^®^ database (https://www.genecards.org) and PubMed. Some of these binding motifs are from studies in Arabidopsis and were not included in the searches.

