## Supplementary figures and images for "Micrococcal nuclease sequencing of porcine sperm suggests a nucleosomal involvement on semen quality and early embryo development"

### Additional file 1

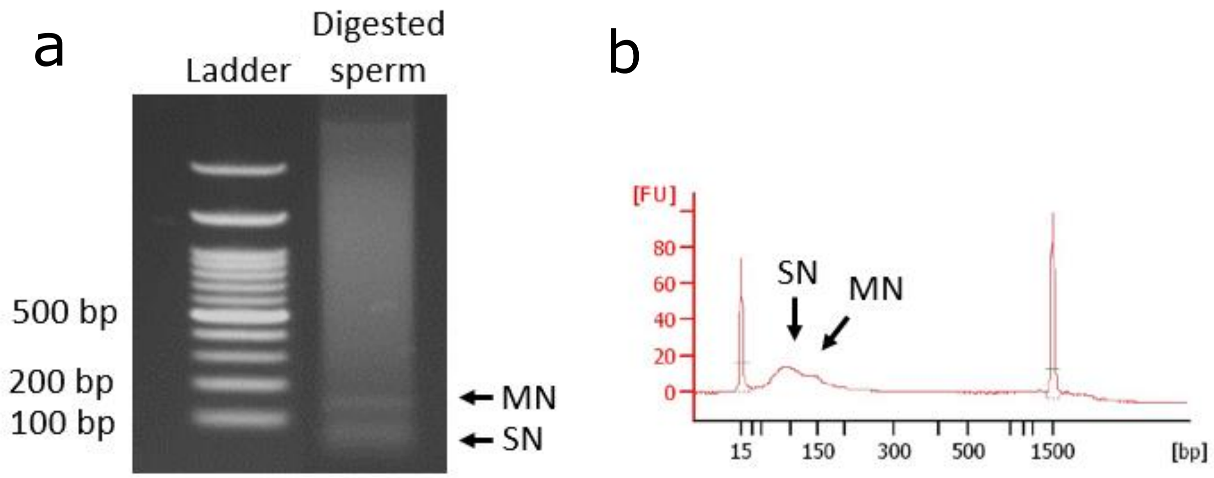

### Additional file 2

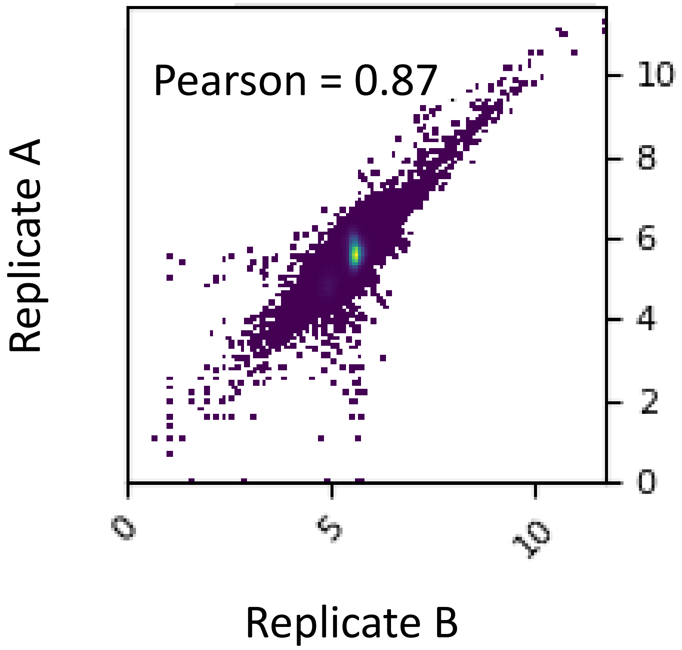
